# A circuit theory of protein structure

**DOI:** 10.1101/023994

**Authors:** G. Sampath

## Abstract

Protein secondary and tertiary structure is modeled as a linear passive analog lumped electrical circuit. Modeling is based on the structural similarity between helix, sheet, turn/loop, and helix pair in proteins and inductor, capacitor, resistor, and transformer in electrical circuits; it includes methods from circuit analysis and synthesis. A ‘protein circuit’ is a one-port with a restrictive circuit topology (for example, the circuit for a secondary structure cannot be a Foster II ladder or a Wheatstone-like bridge). It has a rational positive real impedance function whose pole-zero distribution serves as a compact descriptor of secondary and tertiary structure, which is reminiscent of the Ramachandran plot. Standard circuit analysis methods such as node/loop equations and pole-zero maps may be used to study differences at the secondary and tertiary levels within and across proteins. Pairs of interacting proteins can be modeled as two-ports and studied via transfer functions. Similarly circuit synthesis methods can be used to construct ‘protein circuits’ whose real counterparts may or may not exist. An analysis example shows how a ‘protein circuit’ is constructed for thioredoxin and its pole-zero map obtained. A synthesis example shows how an electrical circuit with a single Brune section is obtained from a specified set of poles and zeros and then mapped to an artificial protein with a helix pair (corresponding to the transformer in the Brune section). Possible applications to folding, drug design, and visualization are indicated.

## 1. Overview

A model of protein structure based on electrical circuits is described. Helices are mapped to inductors, strand pairs to capacitors, turns/loops to resistors, and helix pairs to transformers (coupled inductors). Cys-Cys bonds are capacitors that cause the circuit to fold on itself like the protein modeled. The resulting linear circuit is fully described by its input impedance Z(s), a positive real (p.r.) function of the form P(s)/Q(s), where s is the complex frequency, or equivalently a pole-zero map. The result is a mathematical representation of protein structure with systematic procedures for analysis, synthesis, classification, and design, augmented by an electrical-circuit-based alternative to ribbon diagrams.

The following is a summary of this report. Section 2 gives a brief review of protein structure modeling and a summary of the current approach. Section 3 discusses the derivation of RLCM ‘protein circuits’ from secondary and tertiary structure. Restrictions on ‘protein circuit’ topology resulting from the sequential nature of the protein’s primary sequence are noted. Section 4 looks at the application of circuit analysis methods to ‘protein circuits’ based on impedance functions and to pairs of ‘protein circuits’ using transfer functions. In Section 5, modeling of protein pairs using transfer functions is briefly examined. In Section 6, synthesis methods for the design of protein ‘circuits’ are described. Section 7 concludes with a brief discussion of the potential applications of this approach. An earlier version of this report is available at [1].

## 2. Protein modeling: analytical and synthetic methods

Proteins structure can be considered at three levels: 1) primary, in which a protein is a sequence of amino acids (or equivalently a string of characters drawn from an alphabet of twenty characters); 2) secondary, in which subsequences form three types of geometric shapes: helices, sheets, and turns/loops; and 3) tertiary, in which the secondary structure folds on itself to form complex three-dimensional shapes, within which a number of recognizable ‘motifs’ such as jelly roll, helix pairs, etc. are often seen. One of the main objectives in the study of proteins is to map the primary sequence of a protein to tertiary structure. Also, since form often determines function, knowledge of the relationship of tertiary structure to function is of fundamental importance [2, 3].

The identification of secondary structure consisting of alpha helices, beta sheets, and turns/loops from the primary amino acid sequence of a protein is now fairly routine [4]. In mapping secondary to tertiary structure there are several approaches, including: 1) Analytical methods, which use some kind of minimization of an energy function based on covalent and non-covalent interactions among the side chains and the backbone; some of them are based on lattice models that use cubes [5] or cylinders [6] as structural elements; 2) Synthetic methods, which are aimed at the opposite: deriving a primary sequence that leads to a desired tertiary shape; this reverse process is studied in drug discovery and design and is largely ad hoc [7]; and 3) Visualization studies, which seek to represent graphically the interactions of secondary structure that lead to discernible tertiary substructures seen in classes of naturally occurring proteins [2, 3]; they are often based on diagrammatic representations, such as Richardson’s schematics [2], skeletal structures [3], and TOPS diagrams [8] (which look similar to class diagrams in object-oriented design [9]), and the conventional stick-ball model [2, 3].

In the present work, protein structure is modeled via passive analog lumped electrical circuits [12]. Helix (H), sheet (E), and turn (T) in proteins are mapped to inductor (L), capacitor (C), and resistor (R). By adding capacitive bridges to represent bonds between distant residues and transformers (with mutual inductance M between the coils) to represent helix pairs the resulting RLCM circuit can be used to represent tertiary structure. The equivalence is shown in Figure 1. Standard analysis and synthesis methods [10-14] may then be used to analyze and synthesize ‘protein structures’. As is customary in electrical engineering, the terms ‘circuit’ and ‘network’ are used interchangeably in what follows.

**Figure 1.**
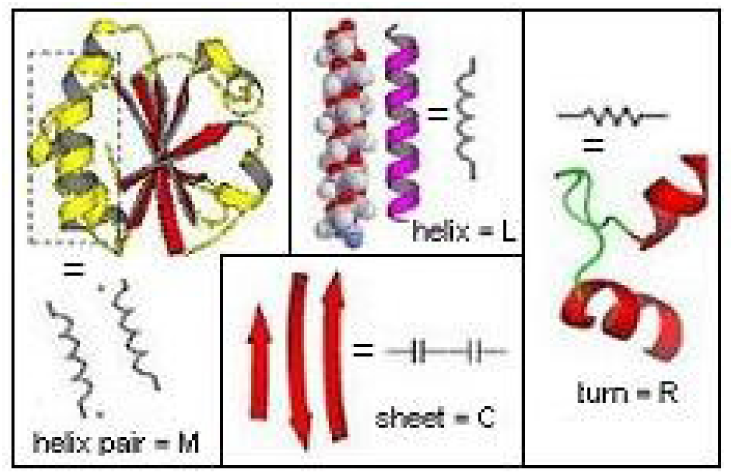
Equivalence of protein secondary elements to R, L, C, and M

## 3. Properties of ‘protein circuits’

### 3.1 Secondary level structure and its circuit analogues

At the secondary level, local chemical constraints and physical forces cause the linear sequence to form helices, strands (which themselves come together to form sheets), and turns that connect strands and/or helices. One helix turn corresponds to about 3.5 residues in the primary sequence, and a sheet has two or more strands. While helices and strands are somewhat rigid, turns are less so (they are thought to flop around loosely in the solvent). Element values are chosen so that every element contributes to the circuit impedance without being swamped out by the others in an appropriate frequency range. A normalization procedure [10] makes the model insensitive to the choice of frequency range. The resulting RLC circuit for secondary structure is named ‘p-RLC-s circuit’. Table 1 shows the mapping from protein secondary elements to electrical circuit elements.

**Table 1.**
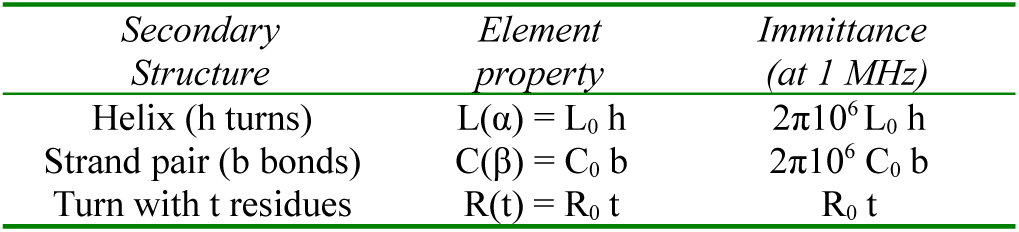
Secondary structure modeling parameters

### 3.2 Tertiary structure

There are two ways in which tertiary structure is obtained from secondary structure:

1. Distant residues in the primary sequence are brought together to be held by a chemical bond, usually a disulphide or a salt bridge [2, 3]. This is modeled as a capacitor between the relevant nodes in the circuit with behavior similar to a strand pair with 1 hydrogen bond (b = 1) leading to a capacitance of C_0_. More realistically, this is multiplied by a constant k_B_ to reflect the strength of the bond or bridge.
2. Helices come together to form a helix pair. This is modeled here as coupled coils with mutual inductance M. The resulting p-RLC-t circuit (with capacitive bridges) or p-RLCM-t circuit (with coupled coils) represents tertiary structure. Tertiary structure modeling is summarized in Table 2.

**Table 2.**
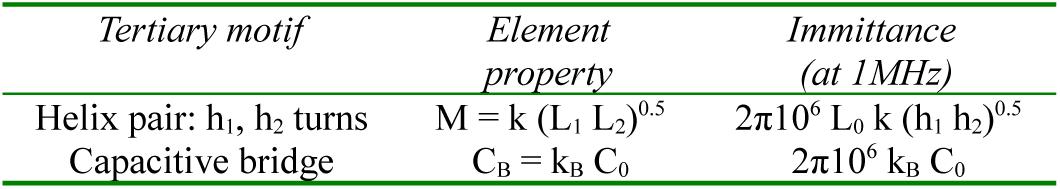
Tertiary structure modeling parameters

*Example: Thioredoxin*

Data from the public domain protein database PDB [15] are used to fix circuit element values, with tertiary elements determined by visual inspection of the ribbon diagram and the schematic for the protein’s entry in PDB. As an example, the p-RLCM-t circuit for thioredoxin (PDB accession id: 1SRX) is shown in Figure 2. Tertiary structure elements are shown as dashed wires (capacitive bridges) or flux linkage lines (between elements of a helix pair) with k (coupling) and M (mutual inductance) values. This manual process can be replaced with a computer program that generates ‘protein circuits’ from PDB data and computes their pole-zero maps using a combination of symbolic and numerical computing [16].

**Figure 2.**
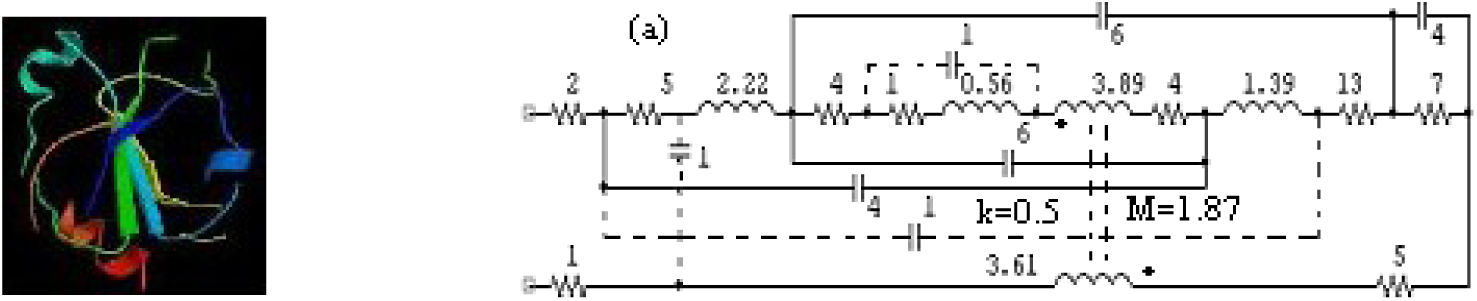
‘Protein circuit’ for thioredoxin (tertiary elements/effects are in dashed lines)

### 3.3 Constraints on ‘protein circuit’ topology

At the lowest level, a protein is a sequence of amino acids held together by a backbone with a characteristic structure. As a consequence, the electrical analogue for secondary structure has an approximate chain or ladder structure (which is modified by the addition of other circuit elements to add more tertiary structure, see below). This places the following fundamental constraints on the circuit topology for the secondary circuit and any other tertiary additions to it:

- an inductor cannot be a shunt element in the ladder
- bridges cannot occur, which also means that the ladder cannot be a series of lattices.

Other consequential constraints are discussed below in the section on synthesis.

### 3.4 The ‘protein circuit’ as a representation device

The circuit diagram of a p-RLC(M) circuit, which is based on a familiar, compact and well-established notation, may be a useful alternative to representations like Robinson’s diagrams or stick figure models [2, 3].

## 4. Circuit analysis methods for ‘protein circuits’

Circuit analysis methods such as node/loop equations and pole-zero maps [12] can be applied to ‘protein circuits’, and the results used to compare compatible characteristics in the two domains as well as classify proteins based on those characteristics. A protein in its primary form is a sequential structure, so that its p-RLC(M) circuit can be viewed as a one-port network that is characterized fully by the input impedance function Z(s).

### 4.1 Circuit properties of a p-RLC(M) circuit

In addition to loop and node equations (‘Kirchhoff’s laws’) there are some other considerations. Thus the protein’s primary sequence has an implicit direction associated with it because of the order in which the protein is synthesized in the cell, which is N-terminal to C-terminal. This is in contrast with passive electrical circuits whose electrical behavior is usually independent of the terminals of the port (with one exception, that of a polarized capacitor, but this behavior is largely a d.c. behavior). It is nevertheless useful to retain the directionality property of the protein’s primary sequence when coupled coils (representing helix pairs) are present since the coupling between them is dependent on the direction of current flow. The dot rule captures [12] this property in a natural way. Thus, in a p-RLCM circuit, the dot is written at the N-end of an inductor in a coupled pair and is used in all circuit computations. The notion of directionality in a passive one-port is therefore not an artificial adjunct.

### 4.2 Input impedance and spectral properties

A linear one-port with lumped R, L, C, and M elements, has an input impedance Z(s) that is a positive real (p. r.) rational function of the complex frequency s. Thus Z(s) = P(s)/Q(s) and is fully specified by its poles (roots of Q) and zeros (roots of P), except for a constant factor. For proteins without any helices or all-helix proteins, the poles and zeros are all on the negative real axis. The corresponding ‘protein circuits’ are RC circuits or RL circuits respectively. In this case, the pole-zero map is easy to compute. For proteins with both helices and sheets, the poles and zeros are in the left half of the s plane and are more difficult to compute symbolically. Several techniques to reduce the effort required are available [16].

When Z(s) is available, the amplitude function |Z(jω)| and the phase function φ(jω) can be computed for s = jω in a routine manner. These three characteristics (pole-zero distribution, amplitude function, phase function), have the potential to act as signatures and/or provide useful classification procedures. In particular the pole-zero map is a two-dimensional descriptor of the ‘protein circuit’; it is reminiscent of the Ramachandran plot [2, 3].

Let N(x) = number of x, where x is a circuit element. Two important properties satisfied by RLCM circuits for proteins are the following:

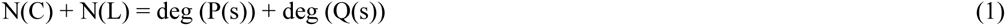

and, when helix pairs are present (corresponding to coupled L’s in the circuit),

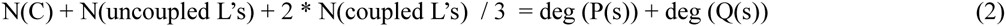

*Example: Properties of circuit model for thioredoxin*

The pole-zero distribution and the phase plot for thioredoxin are shown below for the secondary and tertiary structure circuits. The amplitude response |Z(jω)| has a predictable low-pass behavior and is not shown.

The change in the pole-zero pattern (going from all poles and zeros on the negative real axis for secondary to negative real and some complex poles and zeros for tertiary) can be used to characterize the protein. The phase spectrum can also be examined for signature changes. In the case of thioredoxin, the phase spectrum for tertiary structure has a characteristic humped shape. These change patterns can also be examined for markers that may occur during the folding process.

**Figure 3.**
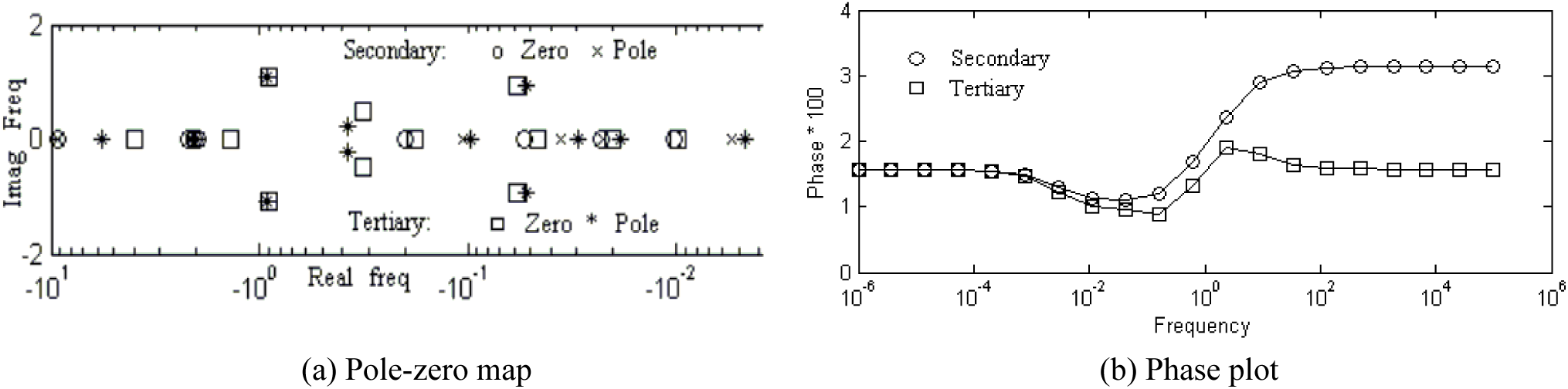
Secondary and tertiary circuit impedance characteristics for Thioredoxin

## 5. Protein pairs: transfer function analysis

A large part of cellular activity can be traced to interacting proteins. Many of the interactions occur because of pairs of proteins coming together (‘docking’) and forming an aggregate shape that causes specific biophysical and/or biochemical reactions to take place. In the model presented here this can be represented by capacitive contacts and/or formation of helix pairs by helices in the two proteins. This essentially results in a two-port network [13, 14] in which one of the ports is represented by the N and C terminals of one of the proteins and the other port by the N and C terminals of the second protein. The protein interaction can be effectively studied through the transfer function T(s) for the two-port circuit.

## 6. Network synthesis methods and an example of ‘protein circuit’ synthesis

An RLCM circuit (one-port or two-port) can be synthesized using frequency domain methods [10, 11, 13, 14] or time-domain-based ones [17]. The last is not as useful in the present context because it is not easy to design circuits with transformers, which means that proteins with helix pairs are excluded. Only frequency domain methods for one-ports are considered here.

A specified positive real Z(s) is implemented with a one-terminal RLCM network. In the first case, standard methods lead to ladder networks that may be canonical or non-canonical in the number of circuit elements used. They include Foster I and II forms, Cauer I and II forms, Brune ladders, Darlington’s method, Bott-Duffin synthesis, and Miyata’s and Kuh’s methods [13, 14]. Different forms can be used in different stages to form mixed ladders, leading to a variety of implementations [12, 14]. However, as mentioned earlier, circuit elements cannot be arbitrarily connected as in a general RLCM network. The following are some additional restrictions:

- A p-RLC-s circuit cannot be a Bott-Duffin type network;
- T-bridges [18] are possible but cannot have an inductor or resistor in the T shunt;
- When turns are present the inductors may sometimes be replaced with lossy ones (L→series LR, RL or LRL, where R represents a turn or loop);
- A-type (but not B-type) Brune sections can be used for tertiary structure with helix pairs;
- In most cases, the dual network does not exist; in particular, non-planar circuits cannot have duals;
- In general, the network realizing a given impedance function is not unique; some of these equivalent networks may or may not correspond to a protein structure.

Starting with an impedance function Z(s) (or equivalently a set of poles and zeros) a secondary structure can be derived and modified to yield a tertiary structure.

*Example of ‘protein circuit’ synthesis*

Consider the impedance function Z(s) = (18s^3^ + 224s^2^ + 457s + 10) / (s^2^ + 4s + 0.06). Following the procedure in [10] leads to the circuit shown in Figure 4(a). It has a single Brune section terminated in a resistor and corresponds to a protein with 68 residues and secondary structure consisting of two α helices, one sheet of two β strands, and three turn-loops (not counting the residues near the terminal ends). A protein shape corresponding to this network is shown in Figure 4(b). The helix pair corresponding to the transformer imposes partial tertiary structure on the protein. More tertiary structure can be introduced as desired through capacitive bridges or by coupling helices subject to physical constraints (such as ‘no knot creation’).

**Figure 4.**
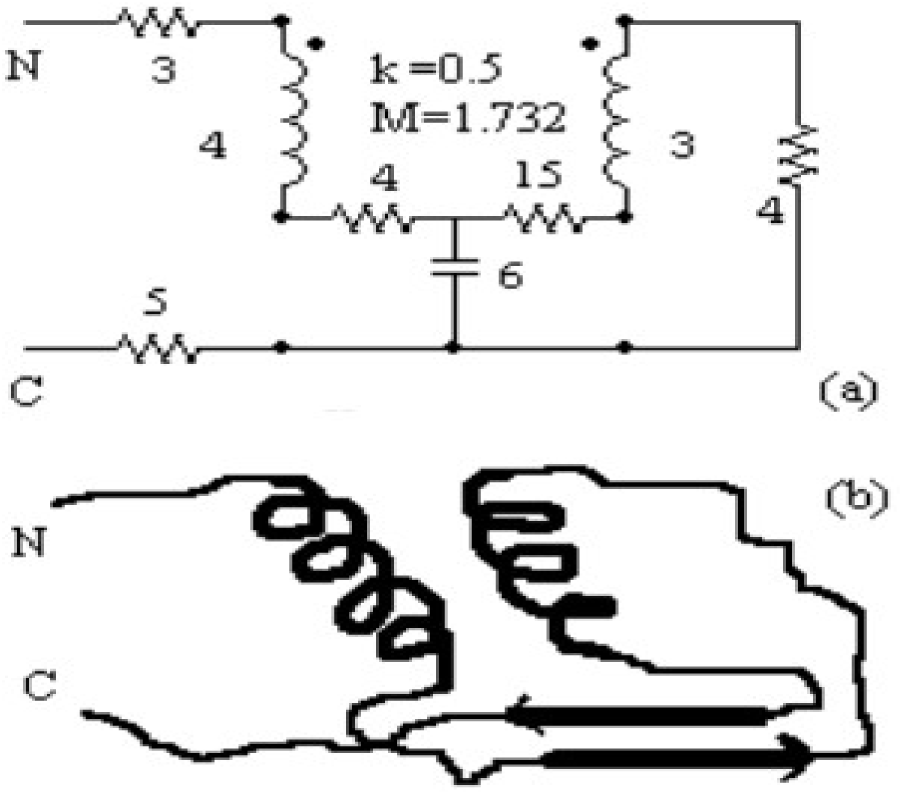
(a) Synthesized circuit for Z(s) = (18s^3^ + 224s^2^ + 457s + 10) / (s^2^ + 4s + 0.06) (b) Corresponding protein shape with secondary and partial tertiary structure (helix pair)

## 7. Discussion

The model presented here provides an electrical-circuit-based alternative to ones based in chemical topology [19] or lattice structures [5, 6]. This approach has several potential uses [1], such as modeling of protein folding (based on sensitivity analysis of the ‘protein circuit’ [10]), searching for proteins that are similar in some sense, drug design and discovery, and the electrical properties of proteins (leading possibly to the use of proteins as nano-level circuits). Circuit simulation provides correlates to chemical structure and behavior of existing proteins, and drug design may be viewed as circuit synthesis using ‘protein circuit’ libraries followed by biochemical synthesis using libraries of designed motifs.

